# TGFβ signaling regulates the response of the skeleton to phosphate

**DOI:** 10.64898/2025.12.01.691659

**Authors:** Emily K. Zhu, Dylan P. Kuennen, Long Tran, Parthena E. Kostalidis, Michael Mannstadt, Lauren E. Surface

## Abstract

Inorganic phosphate (Pi) homeostasis is crucial to organismal health, yet the mechanisms underlying the regulation of it remain unclear. Critically, we lack a clear understanding of the Pi response circuitry in osteogenic cells that identifies altered serum Pi levels and transmits this information to changes in serum FGF23 levels, a key hormone regulating circulating Pi. We utilized genome-wide CRISPR screens in osteogenic Pi-responsive fluorescent reporter cell lines to identify regulators of the response to high phosphate, intersecting those results with loci associated with circulating FGF23 levels by genome-wide association studies (GWAS) and identified a potential role for TGF-β2. We found that each of the three ligands (TGF-β1, 2, 3) can enhance the response to Pi in osteogenic cell lines and ex vivo cultures of calvariae, while inhibitors of TGFβ receptor signaling dampen it. Co-treatment of Pi with TGFβ ligands led to an elevated, synergistic transcriptional induction of *Slc20a1*, which encodes a key Pi importer, which corresponded with an increased intracellular uptake of phosphate. Furthermore, in mice, blocking TGFβ signaling disrupted the induction of FGF23 in mice on a high phosphate diet, resulting in disrupted downstream endocrine control of phosphate homeostasis. Together, these findings reveal a role for TGFβ signaling in the regulation of phosphate homeostasis in osteogenic cells through regulation of cellular phosphate uptake, which in turn contributes to the maintenance of organismal phosphate homeostasis.

## Introduction

Inorganic phosphate (Pi), the simplest soluble form of phosphate and one of the most abundant minerals in the body, plays key roles in metabolism, signal transduction, and nucleic acid synthesis^1^. The regulation of phosphate is crucial for cells as an excess can disrupt cellular reactions, reduce the free energy available for essential processes, and hinder metabolism, among other processes. In the physiological context, high phosphate (hyperphosphatemia) can result in vascular and other extra-skeletal calcifications^2–5^, while its deficiency can result in conditions such as rickets and osteomalacia^6^. Maintaining phosphate homeostasis is therefore fundamental to cellular well-being, with deviations having profound implications for overall health and bone integrity. This maintenance is coordinated by three organ systems: uptake in the intestine, storage in the bone, and excretion through the kidney. Of these, the bone not only functions as a major phosphate reservoir – in mammals, approximately 90% of phosphate is stored in the extracellular matrix of bone tissue^7^ – but also as an active mediator of organismal phosphate homeostasis. A central orchestrator of this inter-organ communication is fibroblast growth factor 23 (FGF23), a hormone primarily secreted by cells of the osteoblast-lineage, particularly osteocytes^8^. FGF23 acts on the kidneys by binding to fibroblast growth factor receptors (FGFRs) in the presence of the co-receptor α-Klotho, resulting in the downregulation of sodium-phosphate transporters NPT2A and NPT2C in the renal proximal tubule, reducing phosphate reabsorption, while also acting to suppress the synthesis of active 1,25-dihydroxyvitamin D, thus altogether lowering serum phosphate levels^8^.

While the effects of FGF23 in maintaining phosphate balance are well-characterized, the mechanisms that govern its production in response to elevated phosphate remain incompletely understood^1,8,9^. Despite being well-characterized in unicellular models such as bacteria and yeast^1,10^, we still don’t understand the cascade that controls how skeletal bone cells detect increased phosphate concentrations and translate this signal into increased FGF23 expression and secretion. Understanding this response mechanism is critical to deciphering the broader regulatory network of phosphate homeostasis.

Some reports have found a role for other organs in the response to changes in levels of circulating phosphate, which then send a signal to the bone to control production of FGF23. For example, kidney glycolysis can function may as a readout of circulating phosphate, with high phosphate resulting in increased release of glycerol-3-phosphate by the kidney, a molecule that can reach the skeleton to stimulate FGF23 production^11,12^. In addition to systemic signals, bone cells themselves likely possess an intrinsic phosphate-sensing capacity to control the release of FGF23 as an endocrine response to high phosphate. For example, elevated extracellular phosphate has been shown to directly activate signaling pathways in osteoblast-lineage cells, including phosphorylation of the FGFR substrate 2α through FGFR and activation of the MEK/ERK cascade, altogether regulating the production and proteolytic cleavage of FGF23^13–23^. As insightfully reviewed in ^8,9^, several studies have identified numerous regulators of this axis, including activation of G_q/11_/PKC signaling^15^, 1,25 dihydroxyvitaminD-mediated activation of the vitamin D receptor^24–28^, and PTH stimulated cAMP generation^29–31^.

Given that control of FGF23 is critical for phosphate homeostasis, it is likely controlled by multiple pathways to ensure a balanced response. To uncover the pathways involved, we took an unbiased, genome-wide CRISPR screening-based approach in osteoblast-lineage cells. Our screens reveal a number of potential regulators of the osteogenic phosphate response. Of these candidates, we show that TGFβ signaling plays a key role in the response to increased extracellular phosphate. We found that TGFβ signaling stimulate the intracellular uptake of phosphate, while enhancing the response to increased extracellular phosphate in cultured osteogenic cells, in *ex vivo* cultures of calvarial punches, and in mice exposed to a high phosphate diet. Our results demonstrate the vital role of osteogenic TGFβ signaling in maintaining organismal phosphate homeostasis.

## Results

To create a genome-wide screen that identifies regulators of the osteogenic phosphate response, we first needed to develop a reporter that could reliably serve as a readout for this response in relevant cells. We utilized the cell lines MC3T3-E1 (a pre-osteoblast-like cell line) and OCY454 (an osteocyte-like cell line) to identify potential reporters, first demonstrating they show robust responses to elevated inorganic sodium phosphate (NaPi). These cell lines were chosen for a) their similarity to osteoblasts and osteocytes *in vivo*, b) their capacity to expand to the numbers needed to undertake a screen, and c) their history in being employed as tools to understand the responses of osteogenic cells to perturbations in mineral ion metabolism. Using qRT-PCR and RNA-Seq, we confirmed that an increase in NaPi induces expression of the expected immediate early genes, including *Fos* and *Egr1,* within 30 minutes, with later induction of mineralization-related genes including *Spp1, Dmp1,* and *Enpp1* in both cell lines (Supp. Fig. 1, Supp. Data 1 & 2), consistent with prior studies on the effects of phosphate on osteogenic cells^19,21,32–35^. (Of note, the basal concentration of the cell culture media used for these experiments is ∼1mM NaPi, similar to the concentration of phosphate in the circulation, to which we added NaPi or NaSO_4_ to the indicated concentrations.) Gene ontology analysis of differentially expressed genes common to both MC3T3-E1 and OCY454 cells treated with elevated NaPi further revealed an enrichment in transcripts related to bone mineralization, tube morphogenesis, enzyme-linked receptor protein signaling, and TGF-β signaling.

**Figure 1.**
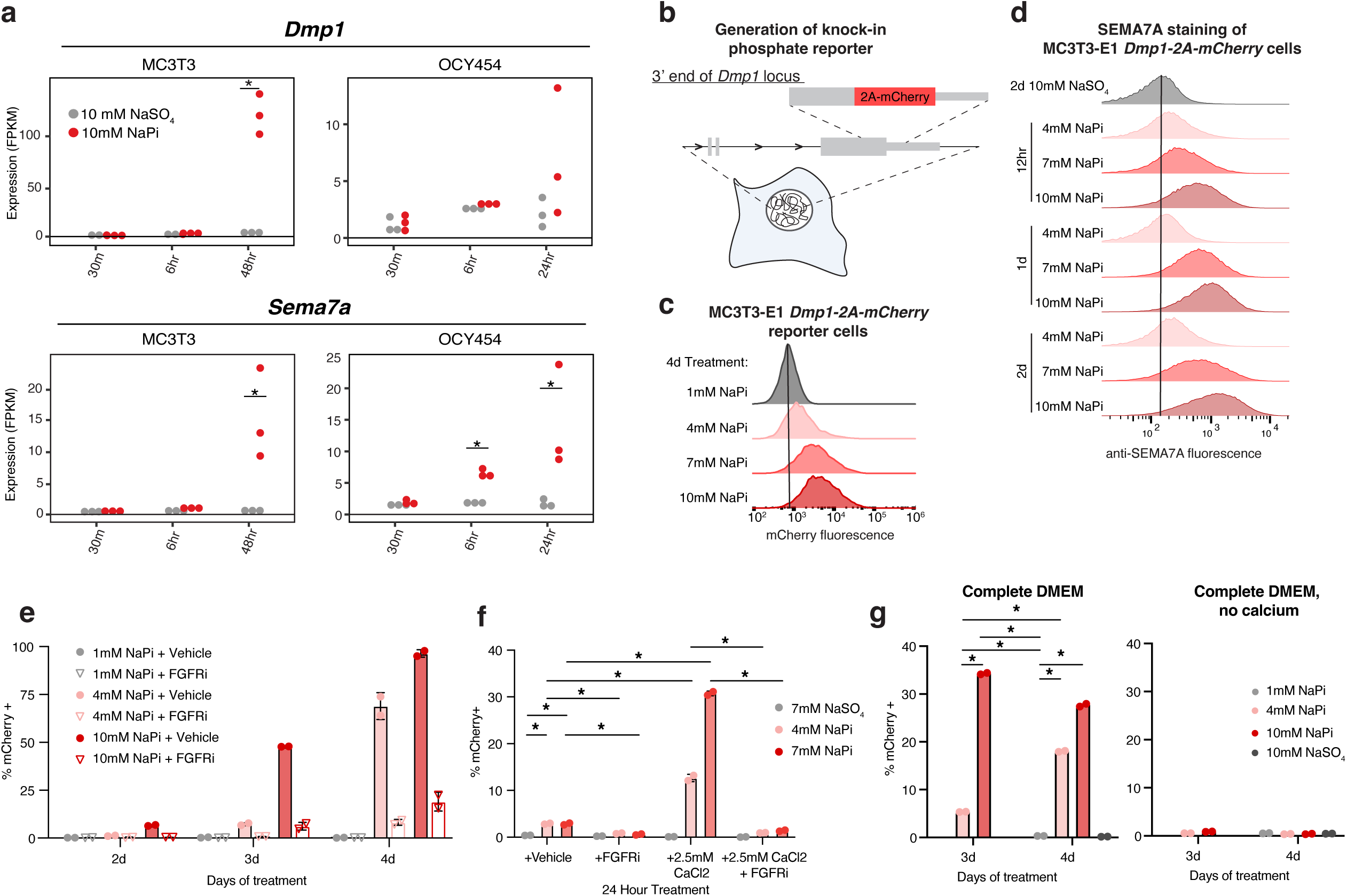
MC3T3-E1 *Dmp1-2A-mCherry* reveal FGF receptor signaling, ERK signaling, and calcium contribute to the osteogenic response to phosphate. **a**. Gene expression as measured by RNAseq of *Dmp1* and *Sema7a* after treatment with 10mM NaPi vs. 10mM NaSO_4_, *: adjusted p-value < 0.05 (from DEseq2). **b**. Schematic of MC3T3-E1 *Dmp1-2A-mCherry* cells, where the endogenous locus of *Dmp1* has a *2A-mCherry* inserted just prior to the stop codon. **c.** Flow cytometry histograms of mCherry fluorescence of MC3T3-E1-Dmp1-2A-mCherry cells exposed to NaPi at indicated concentrations for 3 days. **d.** Flow cytometry histograms of SEMA7A-AlexaFluor488 staining of MC3T3-E1-Dmp1-2A-mCherry cells exposed to NaPi or NaSO_4_ at indicated concentrations and indicated times. **e.** Percent of mCherry+ cells following incubation with 1, 4, or 10mM NaPi + 30 minute pre-treatment with vehicle or 100nM FGFR inhibitor (FGFRi, PD173074) for indicated number of days. **f.** Percent of mCherry+ cells following 1 day incubation with 7mM NaSO_4_, 4mM NaPi, or 7mM NaPi + 30 minute pre-treatment with vehicle, FGFR inhibitor (FGFRi, 100nM PD173074), 2.5mM CaCl_2_, or 2.5mM CaCl_2_ + FGFRi. **g.** Percent of mCherry+ cells following 3 or 4 day incubation in complete DMEM (DMEM + 10% FBS + antibiotics) or complete DMEM, no calcium (No calcium DMEM + 10% FBS + antibiotics) with 1mM NaPi, 4mM NaPi, 10mM NaPi, or 10mM NaSO_4_. n= 2 biological replicates. Significance was assessed by two-way ANOVA, and corrected for multiple comparisons using Bonferroni’s method (*: adjusted p-value<0.05, ns: not significant).

To identify which of the phosphate-responsive genes could be suitable candidates for creating reporters, we focused on genes that were highly upregulated by NaPi, so that flow cytometry could be employed to sort for cells in which these reporters were differentially activated. We chose to focus on two phosphate-induced genes. The first, *Dmp1,* encodes an extracellular matrix protein critical for bone mineralization, and previous work has shown 24-48 hours of high NaPi activates its transcription^19,21,28,34,36^, which we confirmed in our experiments (Fig. 1a, Supp. Fig. 1a). To develop Dmp1 transcription as a fluorescent reporter, using homologous recombination, we inserted a *2A-mCherry* cassette into the 3’ end of the *Dmp1* coding sequence in MC3T3-E1 cells (Fig. 1a). These MC3T3-E1 *Dmp1-2A-mCherry* cells are responsive to NaPi in a dose and time-dependent manner, with fluorescence peaking at 3-4 days of high NaPi treatment (Fig. 1b, Supp. Fig. 2a). For our second reporter, we examined cell surface proteins that get induced by NaPi, in order to utilize antibody-based detection to sort for high and low expressing cells by flow cytometry. This analysis revealed *Sema7a,* which encodes a membrane-bound semaphorin, which gets transcriptionally induced within 6-24 hours of exposure to elevated NaPi (Fig. 1a, Supp. Fig. 1f). Using an antibody against SEMA7A, we observed activation of SEMA7A staining by flow cytometry within 24 hours of high NaPi (Fig. 1c, Supp. Fig. 2b). We reasoned that having these two complementary readouts – one at the transcriptional level (*Dmp1*), the other at the protein level (SEMA7A) – could increase our confidence in the screen’s results and enhance the list of candidate hits.

**Figure 2.**
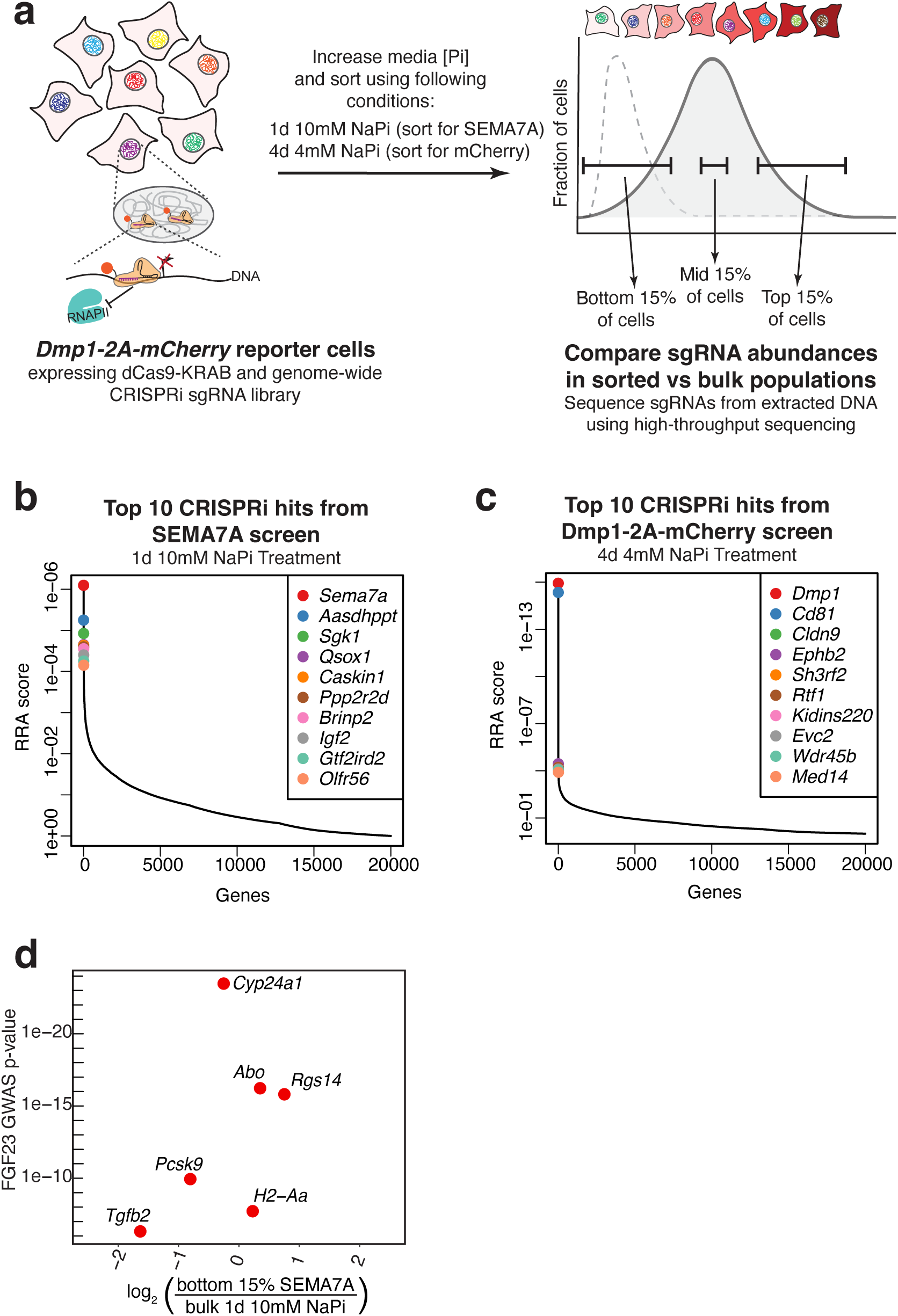
Genome-wide CRISPRi screens reveal regulators of the response to phosphate in osteogenic cells. **a.** Schematic of the genome-wide CRISPR interference screen to identify regulators of the response of MC3T3-E1 cells to phosphate. MC3T3-E1 *Dmp1-2A-mCherry* cells expressing *dCas9-KRAB* are infected with a genome-wide CRISPRi sgRNA library. These cells were exposed to either a) 10mM NaPi for 1 day and stained for SEMA7A for sorting or b) 4mM NaPi for 4 days and sorted by mCherry expression. The bottom 15%, mid 15%, and top 15% of fluorescent cells were sorted. Genomic DNA was then extracted from sorted and bulk populations, sgRNAs were amplified by PCR and then subject to high-throughput sequencing. Dashed gray line represents untreated cells, while solid gray line represents high NaPi treated cells. **b.** Top 10 CRISPRi gene hits from the SEMA7A screen. Data was analyzed using Model-based Analysis of Genome-wide CRISPR-Cas9 Knockout (MAGeCK) software for identification of enriched sgRNAs in biological replicates of the sorted bottom 15% replicates compared to sgRNAs in the bulk unsorted and mid 15% replicates. **c.** Top 10 CRISPRi gene hits from the Dmp1-2A-mCherry screen. Data was analyzed using the MAGeCK software for identification of depleted sgRNAs in biological replicates of the sorted top 15% samples compared to sgRNAs in the bulk, unsorted and mid 15% samples. Corresponding genes were rank-ordered by robust rank aggregation (RRA) scores. The list states the top 10 genes according to RRA scores. **c.** Top 10 CRISPRi gene hits from the SEMA7A screen. Data was analyzed using MAGeCK as in (b), for identification of enriched sgRNAs in biological replicates of the sorted bottom 15% replicates compared to sgRNAs in the bulk unsorted and mid 15% replicates. **d.** Genes associated with FGF23 levels in published GWAS vs. their average enrichment in the SEMA7A CRISPRi replicate screens. Note the mouse gene *H2-Aa* is orthologous to human *HLA-DQA1*.

We then examined whether these reporters were responsive to known regulators of the osteogenic response to NaPi, including FGFR and ERK signaling, and confirmed that co-treatment with inhibitors of FGFR and ERK signaling inhibited the activation of *Dmp1-2A-mCherry* in response to elevated NaPi (Fig. 1d, Supp. Fig. 2c). We next asked whether our reporters respond to calcium, which is known to regulate the phosphate response through the formation of calciprotein particles^37^. We found that co-treatment of elevated NaPi with 2.5mM CaCl_2_ enhanced the activation of the *Dmp1-2A-mCherry* reporter, and that this response was dependent on FGFR signaling (Fig. 1e, Supp. Fig. 2d). Furthermore, using calcium-free media, we showed that calcium is required for this response to NaPi (Fig. 1f). Upon differentiation of our reporter MC3T3-E1 cells into late-stage osteoblasts with beta-glycerophosphate (βGP), we observed activation of SEMA7A in nearly 70% of cells by day 12, and activation of *Dmp1-2A-mCherry* in ∼15% of cells by day 21, which was enhanced by additional NaPi treatment (Supp. Fig. 2e-g). These results suggest that these reporters are primarily responsive to inorganic (NaPi), but not organic phosphate (βGP), as activation of the reporters only occurred following more than a week of βGP treatment compared to the 1 day of NaPi treatment.

Having established that our reporters are responsive to NaPi and to known phosphate regulators, we utilized these cells in genome-wide CRISPR interference (CRISPRi) screens. In these screens, single guide RNAs (sgRNAs) that target promoters genome-wide are introduced to recruit an enzymatically inactive Cas9 (dCas9) fused with a KRAB domain, a transcriptional repressor, to the target promoter to inhibit transcription^38,39^. We introduced the dCas9-KRAB tagged with BFP into our reporter MC3T3-E1 *Dmp1-2A-mCherry* cells, sorted twice for high expression of BFP, and used these polyclonal cells for our genome-wide screens. We tested the efficacy of the dCas9-KRAB in these cells by introducing sgRNAs targeting the promoters of a) *Cd81*, a common control for CRISPRi screens, and b) *Dmp1*, to test a readout of our screen.

Indeed, sgRNAs targeting *Cd81* reduced the binding of fluorescent anti-CD81 antibodies to the cell surface (Supp. Fig. 3a) while sgRNAs targeting *Dmp1* reduced the activation of *Dmp1-2A-mCherry* in response to 3-4 days on high NaPi (Supp. Fig. 3b). These data indicate that dCas9-KRAB effectively inhibited transcription, but that the introduction of dCas9-KRAB didn’t change the phosphate-dependent behavior of our reporter line.

**Figure 3.**
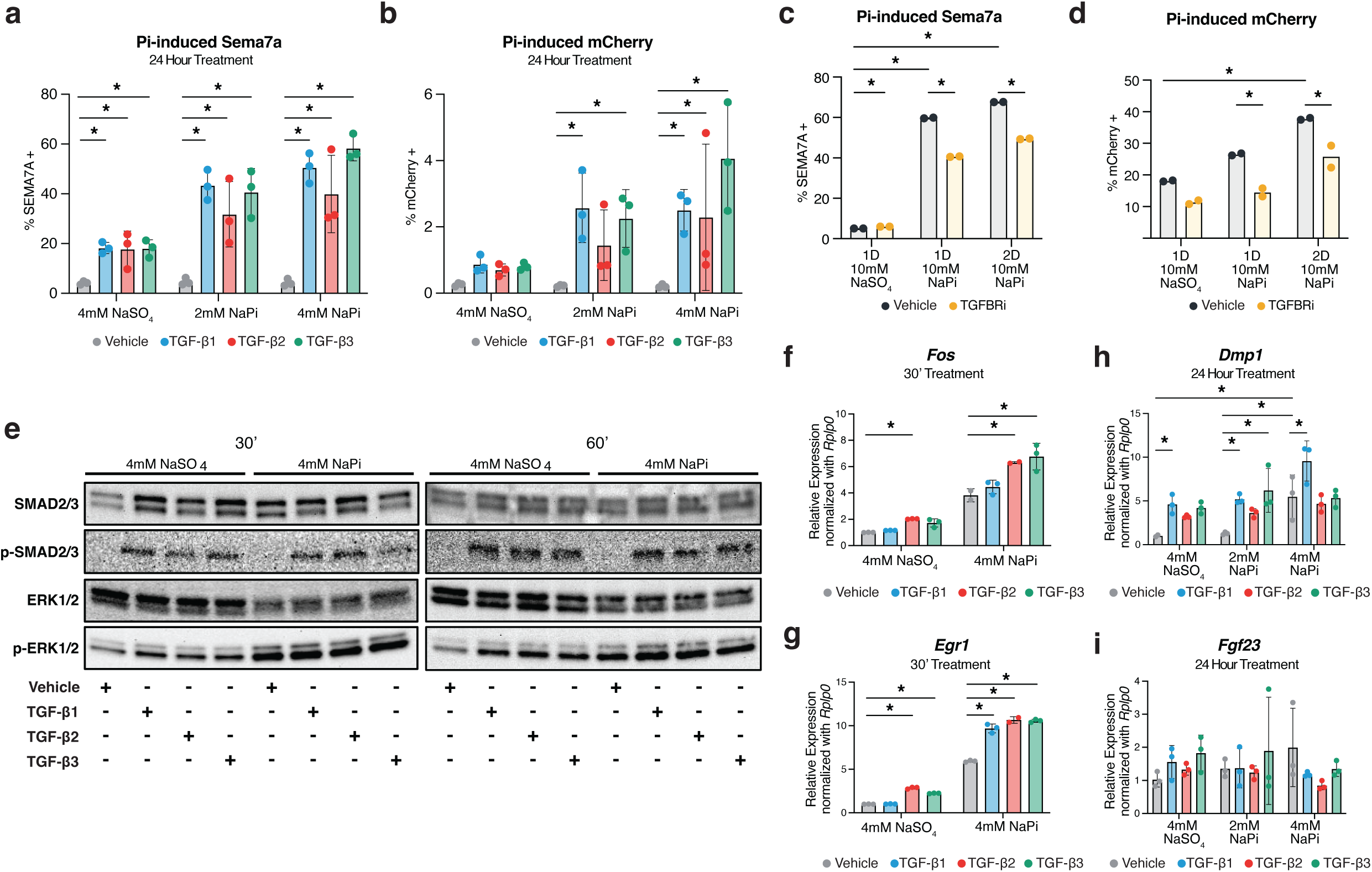
TGFβ signaling enhances the osteogenic phosphate response *in vitro*. **a-b**. Flow cytometry analyses (% positive) of Sema7a and mCherry fluorescence of MC3T3-E1 *Dmp1-2A-mCherry* following incubation with 2mM NaPi, 4mM NaPi or 4mM NaSO_4_, with a 1 hour pre-treatment with either vehicle, TGF-β1, TGF-β2, or TGF-β3 (at 0.4ng/ml). n=3 biological replicates. **c-d.** Flow cytometry analyses (% positive) of Sema7a and mCherry fluorescence of MC3T3-E1 Dmp1-2A-mCherry following incubation with 10mM NaPi or 10mM NaSO_4_, with a 1 hour pre-treatment with either vehicle or 80nM TGFβR inhibitor (TGFBRi, SB-525334) for indicated number of days. **e.** Western blot analyses for SMAD2/3, pSMAD2/3, ERK1/2, pERK1/2 of MC3T3-E1 Dmp1-2A-mCherry cells following 30’-60’ treatment of 4mM NaPi or 4mM NaSO_4_, with a 1 hour pre-treatment of Vehicle, TGF-β1, TGF-β2, or TGF-β3 (at 0.4ng/ml). **f-g.** MC3T3-E1 *Dmp1-2A-mCherry* were treated for 30’ with either 4mM NaPi or 10mM NaSO4 with a 1 hour pre-treatment of either vehicle, TGF-β1, TGF-β2, or TGF-β3 (at 0.4ng/ml). *cFos* and *Egr1* gene expression was assessed by RT-qPCR, with fold change relative to cells treated with vehicle and 10mM NaSO_4_, normalized using expression of *Rplp0*. n=3 technical replicates. **h-i.** OCY454 were treated for 24 hours with 1 hour pre-treatment of either vehicle, TGF-β1, TGF-β2, or TGF-β3 (at 0.4ng/ml). *Dmp1* and *Fgf23* gene expression was assessed by RT-qPCR, with fold change relative to cells treated with vehicle and 10mM NaSO_4_, normalized using expression of *Rplp0*. n=3 biological replicates. Significance was assessed by two-way ANOVA, and corrected for multiple comparisons using Bonferroni’s method (*: adjusted p-value<0.05, ns: not significant).

For our screens, we lentivirally infected our cells with a genome-wide CRISPRi sgRNA library, with 5 sgRNAs per gene at 1000x representation. After selection and minimal propagation, the cells were then split into two conditions: a) treated for 1 day with 10mM NaPi, stained for SEMA7A, and then sorted by percentile of SEMA7A staining (top 15%, mid 15%, and bottom 15%) or b) treated for 4 days with 4mM NaPi, and then sorted by percentile of mCherry expression (top 15%, mid 15%, and bottom 15%) (Fig. 2a). In two biological replicates for each screen, 4.0-7.5 million cells per sorted group and ∼100 million unsorted cells (for comparison to bulk NaPi-treated cells) were collected for gDNA isolation and sequencing. We compared SEMA7A^bottom^ to SEMA7A^mid^ and bulk, unsorted cells using Model-based Analysis of Genome-wide CRISPR-Cas9 Knockout (MAGeCK) analysis software^40,41^ and identified top gene hits (Fig. 2b). Similar analysis for the mCherry screen revealed a distinct, non-overlapping set of top 10 hits. The top hit for each screen was the gene encoding the reporter for that readout (i.e., *Sema7a* was the top hit for the SEMA7A screen and *Dmp1* was the top hit for the Dmp1-2A-mCherry screen). Several top hits were genes previously associated with cellular responses to phosphate, including *Sgk1*, which was shown to regulate vascular calcification of vascular smooth muscle cells in response to NaPi^42–44^ and *Kidins220,* which was recently shown to affect cellular phosphate flux by regulating the activity of the cellular phosphate exporter, XPR1^45–47^, suggesting that the levels of cellular phosphate uptake and efflux impact the responsiveness of osteogenic cells to fluctuating extracellular phosphate concentrations. Furthermore, genes associated with bone mineral density were also identified as top hits, including *Qsox1,* which encodes a protein responsible for regulating the extracellular matrix^48,49^, and *Ephb2,* which encodes a receptor tyrosine kinase associated with regulation of osteogenic differentiation^50–52^. Together, these results suggest that the screens identified genes relevant to the osteogenic response to phosphate.

To move from individual genes to pathways that regulate phosphate homeostasis, we analyzed genes associated with circulating FGF23 levels by genome-wide association studies (GWAS)^53,54^, given the key role of FGF23 in the intestine-bone-kidney axis of homeostatic phosphate control. Of these FGF23 GWAS hits, *Tgfb2,* which encodes the TGFβ ligand TGF-β2, was differentially enriched in our screens (Fig. 2d). Our gene expression analyses of NaPi-treated cells also identified TGFβ signaling as differentially regulated by NaPi (Supp. Fig. 1e), with *Tgfb2* expression decreasing with 24-48 hrs of high NaPi treatment (Supp. Fig. 1f). These observations are consistent with a previous study which showed that TGF-β2 stimulated the transcription of *Fgf23* in osteoblast-like cells^55^. Furthermore, this association between TGFβ signaling and FGF23 has also been observed *in vivo.* In a recent multi-trait analysis of mineral metabolism GWAS (including circulating FGF23 and phosphate), TGFβ signaling was identified as a top regulator effect network^56^, and in conditions where FGF23 is elevated such as chronic kidney disease, TGFβ signaling has also been shown to be elevated^57,58^. We therefore aimed to investigate the role of TGFβ signaling in osteogenic phosphate homeostasis to determine if modulation of this pathway could alter the osteogenic control of phosphate homeostasis.

### TGFβ signaling enhances osteogenic Pi response *in vitro* by increasing intracellular phosphate uptake

The prior association of TGFβ signaling with FGF23 does not necessarily link this pathway with how osteogenic cells are responding to phosphate. To assess the potential roles of TGFβ signaling in this, we examined the effects of the three TGFβ ligands, TGF-β1, TGF-β2, and TGF-β3, on the response to elevations in NaPi. Co-treatment of MC3T3 reporter cells with increasing NaPi concentrations and each TGFβ ligand (0.4 ng/ml) enhanced SEMA7A and DMP1-2A-mCherry, with the strongest induction observed under the highest phosphate levels (Fig. 3a, b). Consistent with this, expression levels of the phosphate-induced genes *Dmp1* and *Enpp1* were also increased by TGF-β2 co-treatment (Supp. Fig. 4a,b). To determine whether TGFβ signaling was required for phosphate’s effects these reporters, we used a TGFβ receptor inhibitor (SB-525334). This inhibitor was tested with a very high phosphate concentration (10mM NaPi) to elicit a robust phosphate response. Inhibition of TGFβ signaling dampened the reponse of both the SEMA7A and Dmp1-2A-mCherry reporters, indicating that the NaPi responses depend on TGFβ (Fig. 3c,d). As cells rapidly respond to increased extracellular phosphate, we next examined intracellular pathways activated by NaPi and TGFβ at early timepoints (30 minutes and 1 hour). Canonically, elevated phosphate induces ERK phosphorylation (pERK), which is required for the induction of downstream responses, including our Dmp1-2A-mCherry reporters (Supp. Fig. 2c)^19,20,33^. If TGFβ signaling intersected the phosphate response pathway upstream of ERK phosphorylation to enhance pERK and its associated downstream effects, this would suggest that ERK phosphorylation would serve as a major signaling hub for the Pi response, integrating signaling from FGFR and TGFβ activation. To assess this, we pre-treated cells with TGFβ ligands followed by 30-60 minutes of 4mM NaPi or NaSO_4_, and found that while TGFβ ligands stimulated SMAD phosphorylation, they did not alter levels of Pi-induced pERK (Fig. 3e). However, gene expression analysis revealed a TGFβ ligand-enhanced upregulation of NaPi-induced immediate early genes *Fos* and *Egr1* as early as 30 minutes post-treatment (Fig. 3f), suggesting TGFβ signaling converges on the Pi response downstream of pERK signaling and upstream of regulation of immediate early gene expression.

**Figure 4.**
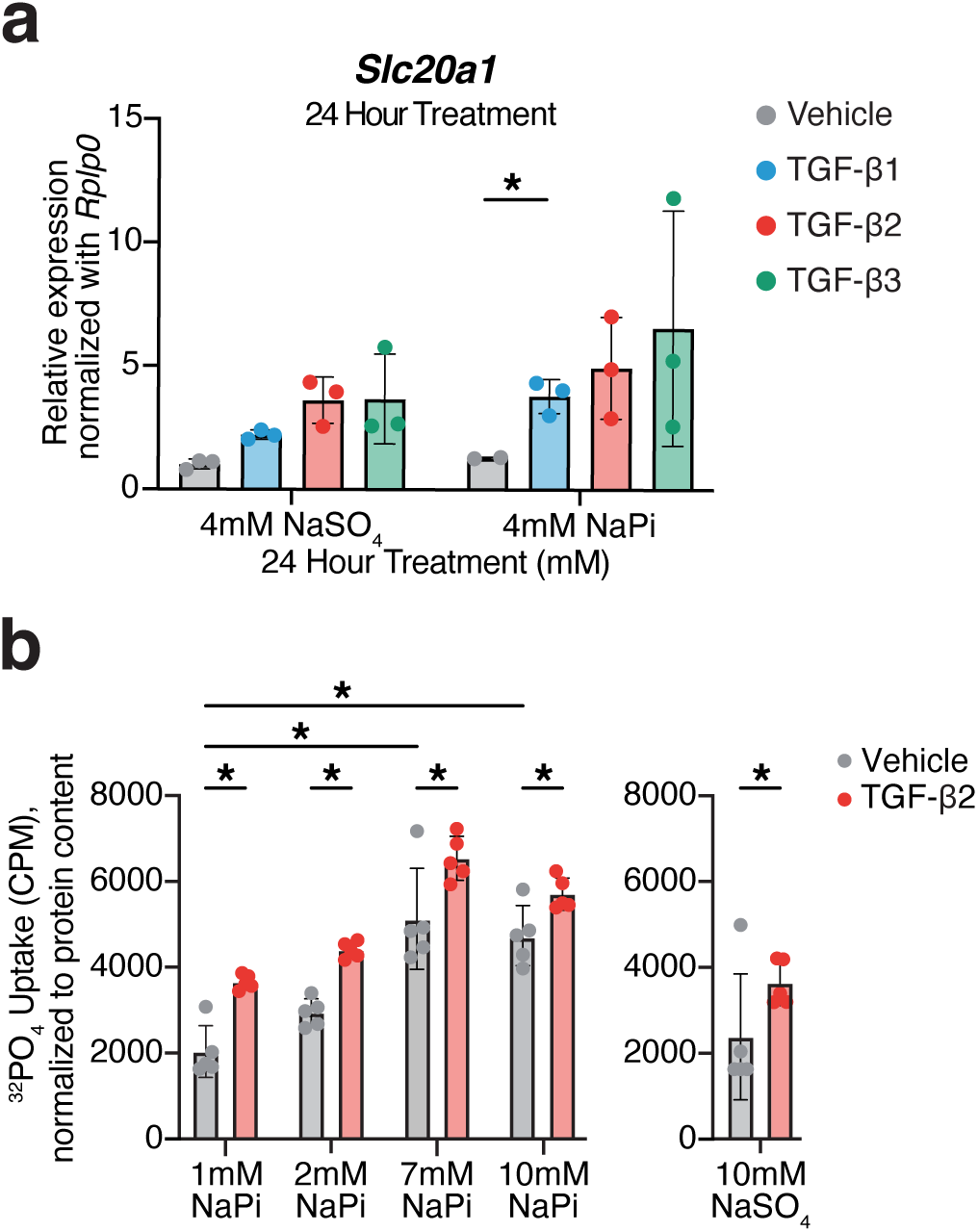
TGFβ signaling stimulates phosphate uptake. **a.** MC3T3-E1 *Dmp1-2A-mCherry* were treated for 24 hours with either 4mM NaPi or 4mM NaSO_4_, with a 1 hour pre-treatment with either vehicle, TGF-β1, TGF-β2, or TGF-β3 (at 0.4ng/ml). *Slc20a1* and *Slc20a2* gene expression was assessed by RT-qPCR, with fold change relative to cells treated with vehicle and 4mM NaSO_4_, normalized using expression of *Rplp0.* n=3 biological replicates. **b.** Cellular phosphate uptake, as measured by incubation of MC3T3-E1 *Dmp1-2A-mCherry* cells in media supplemented with ^32^PO_4_ phosphate for 30 minutes before washing off excess medium and cell lysis. Counts per minute (CPM) were normalized using protein content as determined by PrestoBlue. Prior to incubation with ^32^PO_4_-containing media, cells were treated for 48 hours with indicated NaPi or NaSO_4_ with a 1 hour pre-treatment with vehicle or 0.4ng/ml TGF-β2. n=5 biological replicates. Significance was assessed by two-way ANOVA, and corrected for multiple comparisons using Bonferroni’s method (*: adjusted p-value<0.05, ns: not significant).

To evaluate whether TGFβ influences the osteocytic response to Pi, we utilized OCY454 cells at 14 days of differentiation as a model of osteocyte differentiation. We found *Dmp1* gene expression was stimulated by 24 hour treatment with high NaPi with TGFβ ligand pre-treatment enhancing this induction (Fig. 3h). However, NaPi and TGFβ ligands had largely no effect on *Fgf23* gene expression (Fig. 3i), consistent with prior papers that have not found observed a NaPi-dependent induction of *Fgf23* in osteogenic cells^19,22,23,34^.

However, it remained unclear how TGFβ signaling affects how osteogenic cells sense and respond to phosphate. Prior work in a chondrocyte-like cell line suggested that TGFβ may stimulate intracellular phosphate uptake, by increasing the expression of *Slc20a1*, which encodes one of the two high-affinity, sodium-dependent Pi transporters that mediate the intracellular uptake of Pi^59^. We therefore determined how TGFβ ligands affect the expression of phosphate transporters in bone cells, and found that treatment with NaPi and each TGFβ ligand resulted in an upregulation of *Slc20a1* (Fig. 4a), while expression of *Slc20a2,* which encodes the other sodium-dependent Pi transporter, and *Xpr1*, the main cellular Pi exporter (Giovannini et al., 2013), were largely unaffected by NaPi and TGFβ ligand treatment (Supp. Fig. 4c,d). To assess whether this increased expression of *Slc20a1* in TGFβ treated cells corresponds with increased intracellular phosphate uptake, MC3T3-E1 cells were treated with increasing NaPi concentrations and TGF-β2 (or vehicle) for 48 hours, followed by a 30 minute incubation in ^32^P-orthophosphate containing media. As expected, 48 hour growth in high NaPi media corresponded to increased ^32^P uptake (Fig. 4b). Interestingly, TGF-β2 treatment significantly enhanced this uptake, likely due to the increased expression of *Slc20a1* (Fig. 4a), suggesting TGFβ signaling may be regulating the response to NaPi through upregulation of *Slc20a1* and the promotion of intracellular cellular phosphate uptake.

### TGFβ signaling enhances the response to phosphate in *ex vivo* cultured calvaria

We then utilized an *ex vivo* murine calvarial explant model^60^ to understand how bone cells within their native environment respond to changes in phosphate. We treated calvarial explants from 1 month old wildtype mice with increasing concentrations of NaPi and TGF-β2. As observed in MC3T3 cells, TGF-β2 treatment enhanced expression of *Slc20a1*, as well as *Serpine1*, a canonical TGFβ responsive gene^61^ (Fig. 5b). We then examined the extracellular phosphate concentration in the media, and as expected, phosphate concentrations increased with increasing Pi treatments (Fig. 5c). However, these levels were reduced in TGF-β2 treated calvariae, suggesting TGF-β2 is likely also stimulating Pi uptake in *ex vivo* cultured bones due to increased expression of *Slc20a1*. We then assessed how this affected the expression of *Fgf23*, and found that added NaPi stimulated *Fgf23* expression, and that TGF-β2 co-treatment enhanced this at 4mM NaPi, though not at higher NaPi concentrations (Fig. 5d), changes that are similar to those observed following NaPi only treatment of an osteocytic cell line^24^ and *ex vivo* cultured long bones^22^.

**Figure 5.**
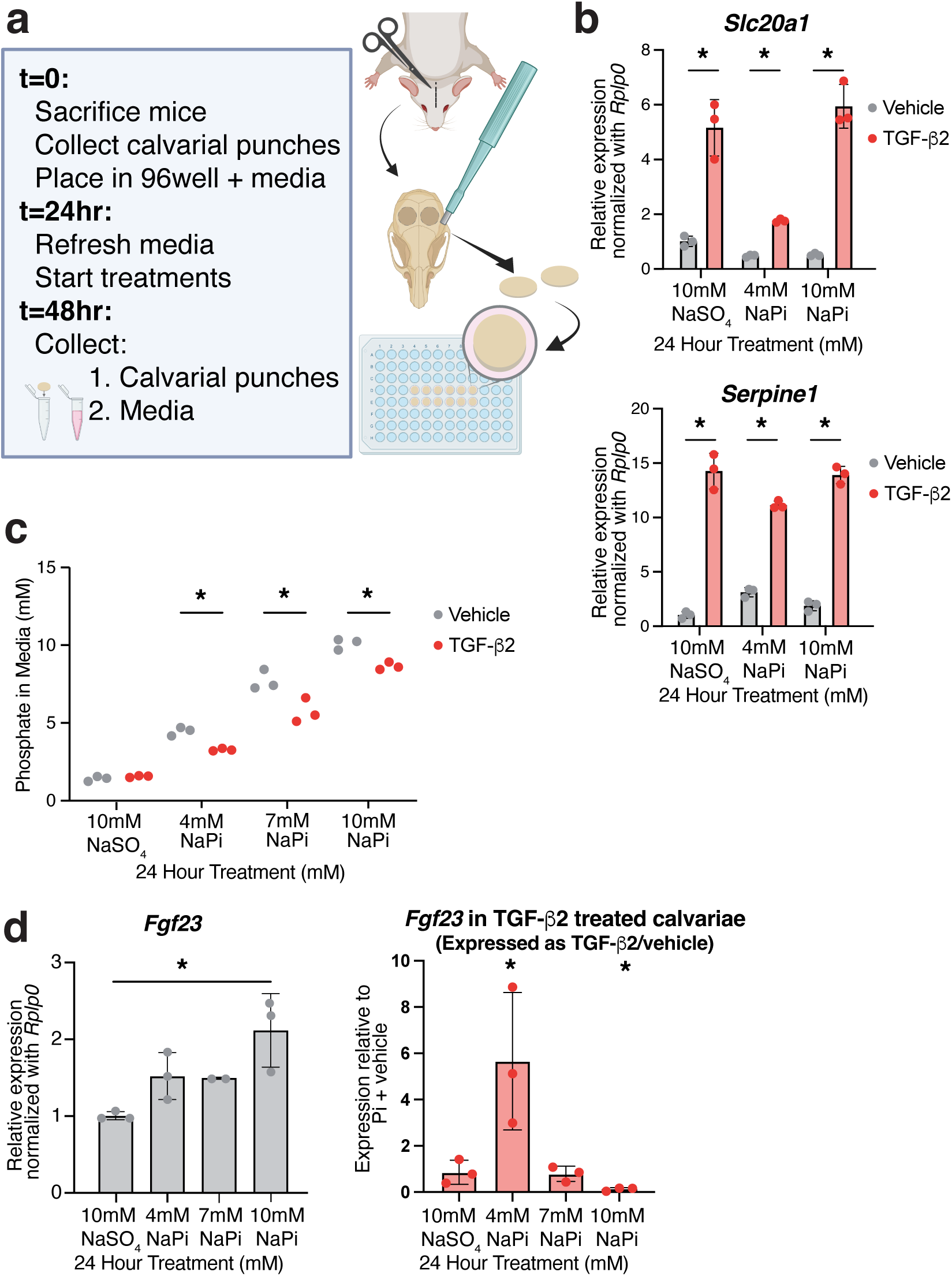
TGFβ signaling enhances the response to Pi in calvarial explant culture. **a.** Schematic of the calvarial explant culture experiments from one month old mice (see Methods for detailed description). Created with Biorender.com. **b.** Gene expression of *Slc20a1* and *Serpine1* as measured by RT-qPCR in calvarial punches treated for 24 hours with indicated NaPi or NaSO_4_ concentrations, with a 1 hour pre-treatment with either vehicle or 0.4ng/ml TGF-β2. Expression is relative to cells treated with vehicle and 10mM NaSO_4_, normalized using expression of *Rplp0.* **c.** Phosphate levels in cell culture media as measured by malachite green assay after 24 hour treatment with indicated NaPi or NaSO_4_, with a 1 hour pre-treatment with either vehicle or 0.4ng/ml TGF-β2. **d.** Gene expression of *Fgf23*. Left graph is *Fgf23* expression with indicated NaPi or NaSO_4_ concentrations alone. Right graph is *Fgf23* expression in TGF-β2 treated cells compared to vehicle treated cells at that same NaPi or NaSO_4_ concentration. n=3 biological replicates for all experiments. Significance was assessed by one-way ANOVA, and corrected for multiple comparisons using Bonferroni’s method (*: adjusted p-value<0.05).

### Blocking TGFβ signaling disrupts circulating iFGF23 *in vivo*

We next examined the *in vivo* impact of TGFβ signaling on FGF23 production in response to high phosphate. Wild-type C57BL/6 male mice were given a control or high phosphate diet for one week (0.6% Pi vs. 1.8% Pi w/w) and injected three times over the course of the week with either a pan anti-TGFβ neutralizing antibody (1D11, 5mg/kg) or control IgG. As expected, antibody treatment reduced circulating serum TGF-β2. On the high phosphate diet, IgG treated mice exhibited the expected increase in circulating intact FGF23 (iFGF23), whereas antibody-treated mice displayed a blunted endocrine response with reduced iFGF23 as well as decreased femoral *Fgf23* expression (Fig. 6a,b). While femoral *Tgfb1* and *Tgfb2* showed a trend toward increased expression in antibody treated mice, this was not significant (Fig. 6b), and may reflect local compensation for the reduced TGFβ signaling.

**Figure 6.**
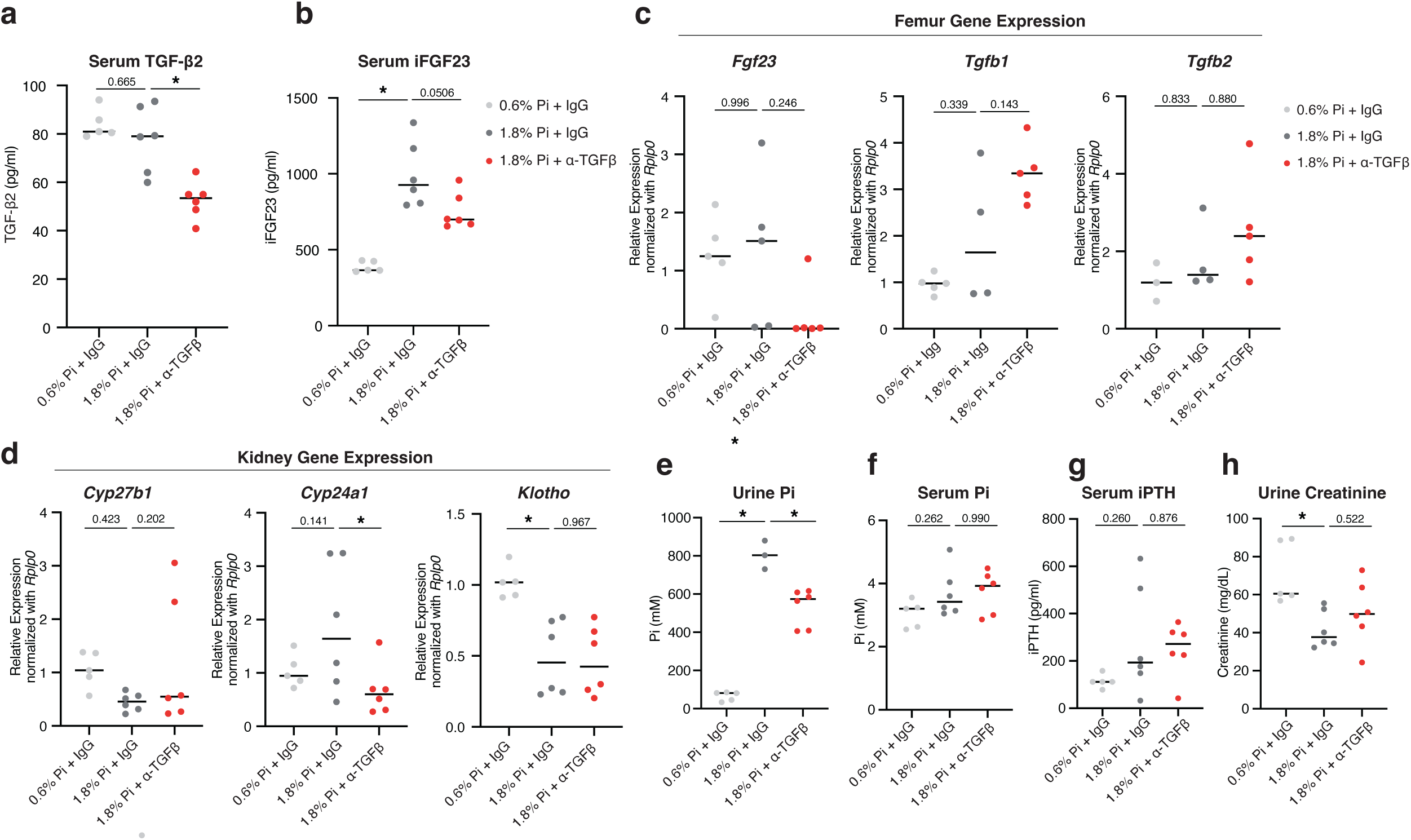
TGFβ signaling regulates release of FGF23 in response to a high Pi diet *in vivo*. 11 week old wildtype mice were treated with either control IgG or neutralizing pan-TGFB antibody (1D11) (at 5mg/kg) three times over one week and were placed on a normal phosphate (0.6% Pi) or high phosphate (1.8% Pi) diet. **a.** Serum TGFB2 and **b.** serum iFGF23 were assessed by ELISA. **c.** Femoral *Fgf23*, *Tgfb1*, and *Tgfb2* gene expression was assessed by RT-qPCR, with fold change relative to mice on a normal phosphate diet treated with IgG, normalized using expression of Rplp0. **d.** Kidney *Cyp27b1*, *Cyp24a1*, and *Klotho* gene expression was assessed by RT-qPCR, with fold change relative to mice on a normal phosphate diet treated with IgG, normalized using expression of Rplp0. **e.** Urine phosphate measured was assessed with colorimetric quantification. (n=3 for mice on 1.8% Pi diet treated with IgG due to loss of samples). **f.** Serum phosphate was assessed with colorimetric quantification. **g.** Serum iPTH was assessed by ELISA. **h.** Serum creatinine was assessed by ELISA. n= 4-6 biological replicates for all experiments. Significance was assessed by two-way ANOVA, and corrected for multiple comparisons using Šidák’s correction (*: adjusted p-value<0.05, ns: not significant).

FGF23 suppresses synthesis of active 1,25 dihydroxyvitaminD (1,25D) in the kidney through reciprocal control of the expression of *Cyp27b1,* which encodes the enzymes that catalyzes formation of 1,25D, and *Cyp24a1,* the enzymes that catalyzes catabolism of 1,25D. In IgG treated mice, as expected, the high phosphate diet and associated increase in iFGF23 tended to suppress *Cyp27b1* and induce *Cyp24a1* expression. Compared with IgG treated mice, anti-TGFβ antibody-treated mice on a high phosphate diet had largely unchanged *Cyp27b1* but significantly decreased *Cyp24a1* expression as compared to control IgG treated mice, suggestive of impaired control of renal Vitamin D synthesis (Fig. 6c), consistent with blunted iFGF23 in antibody-treated mice. Renal *Klotho* expression, which is necessary for FGF23 activity in the kidney^62,63^, decreased in response to the high Pi diet, but remained unaffected by the antibody treatment (Fig. 6c).

Normally, under high phosphate conditions, FGF23 acts on the kidney to limit phosphate reabsorption, leading to increased urinary phosphate excretion^8^. Consistent with this, mice on a high phosphate diet showed elevated urinary phosphate. However, antibody-treated mice excreted less urinary phosphate (Fig. 6d) and a slight, though insignificant rise in serum phosphate (Fig. 6e), suggesting that the blunted FGF23 limits the kidney’s ability to clear excess phosphate while on a high phosphate diet, consistent with disrupted endocrine compensation and renal homeostasis.

## Discussion

Our screens took an unbiased approach to assess the potential regulators of this osteogenic phosphate response and identified a multitude of genes that differentially regulate this response. The addition of TGFβ enhanced the phosphate response in osteogenic cells and calvarial explant cultures, likely through increasing intracellular phosphate uptake. Expressed across nearly all tissues in the body, transforming growth factor-beta signaling regulates a wide range of cellular and physiological processes^64,65^. Owing to its broad physiological influence, targeting TGFβ signaling has emerged as a promising therapeutic modality for several conditions, including cancer^66,67^, osteogenesis imperfecta^68,69^, and inflammatory bowel disease^70^. Stored in the bone matrix and released during bone resorption by osteoclasts, TGFβ has well-documented roles in skeletal development as reviewed in ^71^ and bone remodeling, including through regulating osteoclast and osteocyte functions^72–75^. Our findings add another layer to this and suggest that TGFβ’s influence extends beyond development and structural remodeling to include effects on mineral metabolism and endocrine maintenance of homeostasis.

Indeed, our results suggest that TGFβ signaling may impact response to phosphate through increasing intracellular phosphate uptake, consistent with work in chrondrocytes ^59^. Other hits from our screens also suggest that cellular phosphate uptake and flux may be important in regulating the osteogenic response to Pi, including *Kidins220*, which encodes a protein that affects cellular phosphate flux by regulating the cellular phosphate exporter XPR1^45–47^. Previous work has established that extracellular phosphate affects cellular functions across a range of mammalian and that mammalian cells are able to regulate their uptake and efflux in response to these changing conditions^46,76,77^ , as we have observed in MC3T3-E1 cells (Fig. 4b). It appears that altered phosphate flux may provide cell type specific functions^46,78–82^ , and mutations in transporter genes (including *Xpr1, Slc20a1, Slc20a2*) are associated with differential disease patterns^1^ . With regards to the role of phosphate in regulating the production of FGF23 in osteogenic cells, prior work suggested that the phosphate importer SLC20A2 was required for phosphate-mediated FGF23 secretion^22^, and that this may work primarily through heterodimerization of SLC20A1 and A2 and not by SLC20A1/A2-mediated phosphate uptake directly^33^. Altogether, it remains unclear how intracellular phosphate flux and XPR1-mediated phosphate export may regulate FGF23 production. Future studies on the roles of phosphate import and export machinery in osteogenic cells could shed light the intersection between phosphate flux, TGFβ, and the regulation of FGF23.

There are many diseases where phosphate homeostasis and control of FGF23 is broken, such as chronic kidney disease (CKD)-associated hyperphosphatemia, which is accompanied by highly elevated FGF23. This FGF23 elevation is associated with several pathological disorders, and in CKD, FGF23 has been independently associated with an increased risk for mortality, through increased cardiovascular events, which is the leading cause of death for CKD patients^83,84^, and it is thought that elevated FGF23 may be targeting cardiomyocytes to induce cardiac hypertrophy^85,86^. Our study suggests that it would be worth exploring the roles for TGFβ in these conditions. Indeed, a mouse model of chronic kidney disease treated with the same anti-TGFβ neutralizing antibody found a reduction in FGF23 after 4 weeks of treatment^58^ . Since TGFβ signaling is amenable to perturbation through previously developed therapeutics for other conditions, it could potentially offer therapeutic avenues for both phosphate-related diseases and complications from FGF23 overload.

A potential caveat to this is that our *in vivo* work used a pan anti-TGFβ neutralizing antibody to reduce FGF23 levels. Because this is not specifically targeted to the bone, we can’t determine whether the effects on FGF23 are direct or indirect, for example, by targeting phosphate transport elsewhere in the body. Future experiments to tease apart the role of bone-specific TGFβ signaling will be required. This could include employing bone-targeted anti-TGFβ antibodies, as have been trialed with a bone-targeted anti-sclerostin antibody^87^, or utilizing mice with an osteocyte-specific defect in TGFβ signaling^73^. While anti-TGFβ antibodies have demonstrated promise in reducing FGF23 in mouse models of CKD^58^, it remains unclear how to interpret those results, and perhaps a bone-targeted version could help distinguish effects of anti-TGFβ on skeletal vs. extraskeletal tissues.

Overall, our study demonstrates how genome-wide CRISPRi screens, when coupled with relevant reporters and physiological *ex vivo* models, can be used to illuminate mechanisms of phosphate homeostasis, generating directions that can be followed up with *in vivo* models. In this study we focused on TGFβ signaling, but our screens also point to other pathways that could be promising for future studies of organismal phosphate homeostasis, including other extracellular matrix proteins including QSOX1 and downstream signaling through SGK1.

Furthermore, while this study focused on the kidney-bone axis, the increasing availability of organoid and *ex vivo* culture models from a variety of tissues makes this approach applicable to investigate other cases of inter-organ communication and maintenance of endocrine homeostasis.

## Materials & Methods

### Cell Culture

All procedures related to cell culture were performed within a sterile Class II cell culture hood. Cells were cultivated in standard plastic culture-treated dishes and maintained in an incubator at 5% CO_2_ before undergoing experimental treatments. All reagents were both sterile and pre-warmed to 37°C. We obtained the MC3T3-E1 clone 14 cell line, a calvarial pre-osteoblast cell line, from ATCC (American Type Culture Collection). These cells were cultured in Minimum Essential Media α (MEM α) without ascorbic acid (Gibco, Thermo-Fisher Scientific), supplemented with 10% Fetal Clone III (FCIII, HyClone), and 1X Antibiotic-Antimycotic (Invitrogen, Thermo-Fisher Scientific). The OCY454-12H cell line are an osteocyte cell line kindly provided by Dr. Paola Divieti Pajevic (Boston University). These cells were cultured in Minimum Essential Media α (MEM α) 1X with ascorbic acid (Gibco, Thermo-Fisher Scientific), supplemented with 10% Fetal Clone III (FCIII, HyClone), and 1X Antibiotic-Antimycotic (Invitrogen, Thermo-Fisher Scientific). MC3T3-E1 cells were seeded at a density of 25,000 cells per cm^2^ of well surface area and incubated at 37°C. OCY454-12H cells expanded in 33°C and when ready for experimentation, seeded at a density of 50,000 cells/ml, incubated for 48 hours to allow for cells to adhere to the plate, then moved to 37°C for differentiation.

### RNAseq

RNA from treated cells was extracted with RNeasy kit (Qiagen). For RNASeq and bioinformatics analysis (BGI Americas), sequencing was performed on the BGISEQ platform (single end 50bp reads) platform. Bioinformatics analysis was performed on QC, cleaned reads by mapping reads using Hisat v2.0.5 to the Mus musculus genome, and the DESeq2 algorithm was employed for differential expression analysis (Supp. Tables 1&2). Gene enrichment analysis of significantly upregulated or downregulated genes was performed with Metascape^88^ (http://metascape.org).

### Generating DMP1-2A-mCherry expressing MC3T3-E1 cells CRISPRi Screen

MAGeCK (version 0.5.9) was used for screen hit identification, quality control, and visualization. The NGS data was analyzed using the ‘count’ and ‘test’ commands from MAGeCK with default parameters and zero count removal. A list of mice housekeeping genes were used as a negative biological control. MAGeCK R markdown, a visualization module of MAGeCK, containing the result of the analysis was run in R to generate the illustrations.

### Sodium phosphate, small molecule, and ligand treatments

For all NaPi treatments, the noted concentrations are the final concentration in the media. The base media (MEM α without ascorbic acid) contains ∼1mM NaPi, and to then reach the final concentration noted we add NaPi pH7.4 (0.28 M NaH_2_PO_4_, 0.717 M Na_2_HPO_4_) to reach that final concentration, e.g. for 4 or 10mM NaPi, we added 3 or 9 mM NaPi.

Small molecule inhibitors (FGFR1 inhibitor, PD173074, 100nM, Cayman; TGFβRI inhibitor, SB525334, 80nM, Selleck Chemicals; MEK inhibitor, U0126, 10μM, Cell Signaling Technologies) or vehicle controls were added 30 minutes prior to addition of added NaPi. For treatment with TGF-β ligands, cells underwent a 30 minute or 1 hour pre-treatment in duplicate or triplicate with either a vehicle control (4mM HCl + 0.1% BSA) or 0.4 ng/ml recombinant mouse TGF-β1, TGF-β2, or TGF-β3 protein (TGF-β1, R&D Systems; TGF-β2, R&D Systems; TGF-β3, Sigma-Aldrich). After the pre-treatment, sodium phosphate (NaPi pH7.4) or ionic control NaSO_4_ was added at indicated concentrations and times. For treatments longer than 24 hours, media and treatments were refreshed every 24 hours. Following the treatment period, cells were harvested for flow cytometry, RNA, and protein analysis, as detailed below.

### Immunoblotting

Cell cultures were lysed using 100 μL Pierce RIPA Buffer (ThermoFisher Scientific) per 10 cm^2^ dish supplemented with protease inhibitors (Roche). Samples were then sonicated for approximately 10s three times and stored on ice. Total cell lysate protein concentrations were determined using the Pierce Bicinchronic Acid (BCA) Protein Assay kit (ThermoFisher Scientific) according to the specifications detailed in the manufacturer’s protocol. Western blot analysis was performed using 30 μg protein of cellular lysates and run on a Novex WedgeWell 4-20% Tris-Glycine Gel (Invitrogen, ThermoFisher Scientific) for one hour at 100mV. Protein was transferred to Immobilon-E PVDF transfer membrane for 75 minutes at 100V. The protein transfer efficiency was assessed using Ponceau Red to demarcate total protein. The membrane was washed several times in distilled H_2_O and again in 1X tris buffered saline with 0.1% Tween 20 detergent (TBST). Primary and secondary antibodies were diluted in 1X TBST with 5% BSA unless otherwise noted. To block primary antibody binding to non-specific protein, the membrane was incubated in 1X TBST with 5% bovine serum albumin (BSA) for one hour at room temperature. Membranes were incubated overnight with primary antibody dilutions at 4°C with light rocking with SMAD2/3 (#8685, Cell Signaling Technology, diluted at 1:1000),

Phospho-SMAD2/3 (#8828, Cell Signaling Technology, diluted at 1:1000 in 1X TBST with 5% w/v nonfat dry milk), ERK1/2 (#4695, Cell Signaling Technology, diluted at 1:1000), or Phospho-ERK1/2 (#9101, Cell Signaling Technology, diluted at 1:1000). The next morning, primary antibodies were removed and the blots were washed in 1X TBS with 0.1% Tween (TBST) solution three times. Secondary antibody Anti-rabbit IgG (#7074, Cell Signaling Technology, diluted at 1:1000), was incubated at room temperature with light rocking. The membrane was washed twice more with TBST and detection was performed using the Pierce ECL Western Blotting Substrate (ThermoFisher Scientific) according to the manufacturer’s recommendations in conjunction with the ChemiDoc imager (Bio-Rad).

### Flow cytometry analysis

MC3T3-E1 *Dmp1-2A-mCherry* cells were washed once with PBS and dissociated from culture plates using TrypLE. Cells for each experimental condition were then collected and centrifuged at 1000 rpm for 5 minutes. Supernatant was removed and cells were then washed three times with ice cold 1X PBS + 1% BSA. Cells incubated for 1hr at 4°C with anti-SEMA7A (2.5μg/ml, AF1835, R&D Systems) or goat IgG (AB-108, R&D Systems) in 1xPBS + 1% BSA and gently mixed every 15 minutes. Cells were then washed three times more with ice cold 1X PBS + 1% BSA. Cells were then incubated in secondary antibody (2μg/ml, anti-goat AlexaFluor-488, ab150129, Abcam) in 1X PBS + 1% BSA. After washing three times with 1X PBS + 1% BSA, cells were analyzed using an Attune NxT flow cytometer with a CytKick Autosampler (ThermoFisher Scientific) using both the blue and yellow lasers. Analysis of flow cytometry data utilized the percent positive value of single cells gated for fluorophore intensity.

### Gene expression analysis

For homogenization, cells were suspended in 500 μl of Trizol reagent (Invitrogen, ThermoFisher Scientific); tissue samples were pulverized in liquid nitrogen and suspended in 1000 μl of Trizol reagent. Samples were either stored at -80°C or immediately purified in accordance with the manufacturer’s protocol. RNA was extracted using TRIzol (Invitrogen, ThermoFisher Scientific) in accordance with the manufacturer’s protocol, an additional chloroform and additional ethanol wash was used for tissue samples to increase purity. Total RNA and RNA purity were assessed using a NanoDrop One Microvolume UV-Vis Spectrophotometer (ThermoFisher Scientific). To generate complementary DNA (cDNA), samples were processed using the SuperScript IV Reverse Transcriptase system (ThermoFisher Scientific) following the manufacturer’s specifications. The resulting cDNA samples were subject to qRT-PCR using primers designed using NCBI’s Primer-blast software to span exon-exon junctions and purchased from Integrated DNA Technologies (IDT) (primer sequences in Table 1). Ribosomal protein lateral stalk subunit P0 (*Rplp0*) served as an endogenous control. qRT-PCR was run with triplicate reactions using the PowerUp SYBR Green Mastermix (Applied Biosystems, ThermoFisher Scientific) on a QuantStudio 5 or Power SYBR Green PCR Mastermix (Applied Biosystems, ThermoFisher Scientific) on a QuantStudio 6 Real-Time PCR System (Applied Biosystems, ThermoFisher Scientific) following the manufacturer’s recommendations for cycling conditions. Gene expression was quantified relative to *Rplp0* levels using the 2-ΔΔC_T_ method, with the control sample noted in each relevant figure.

Forward and reverse primer sequences:

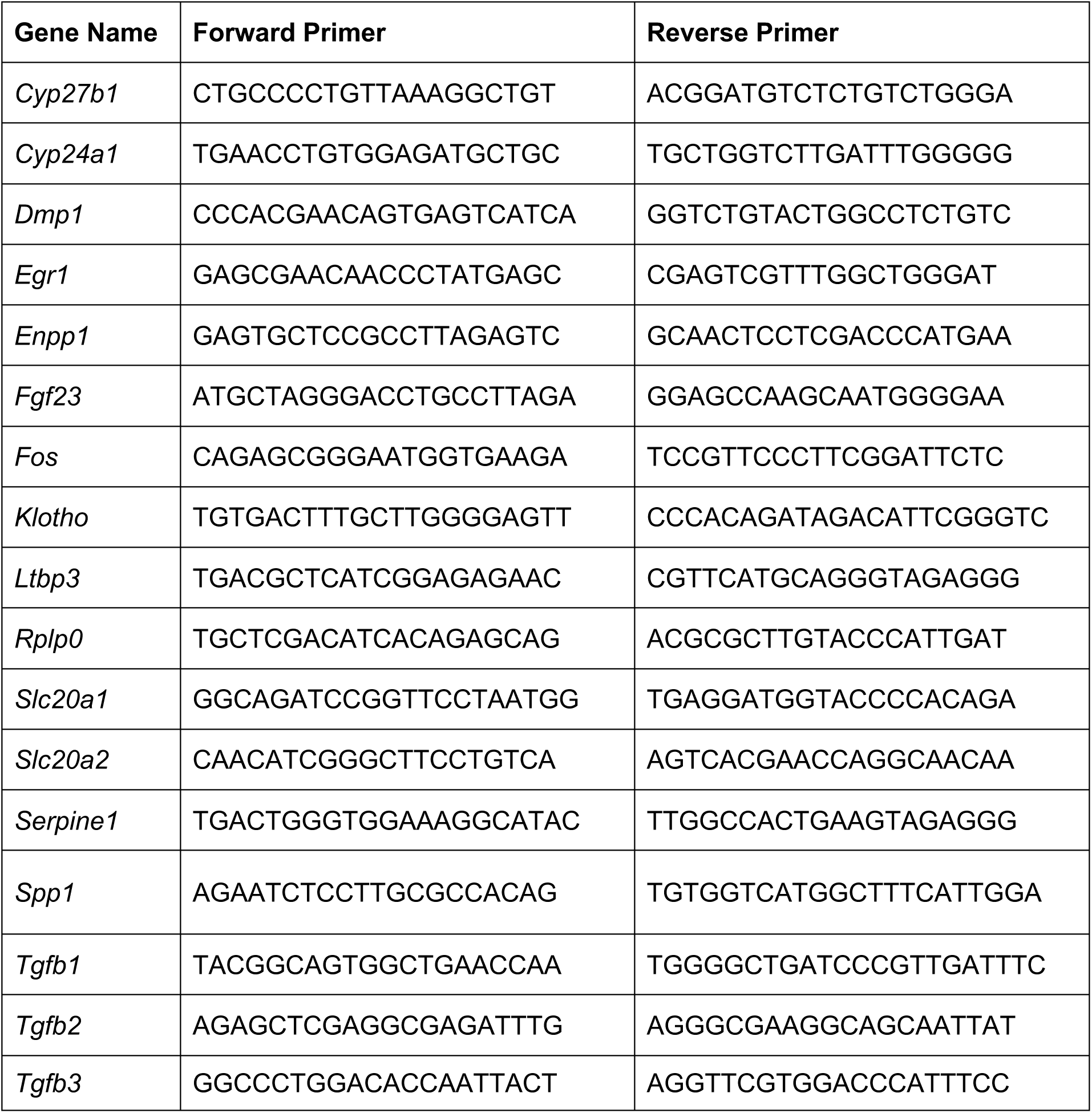
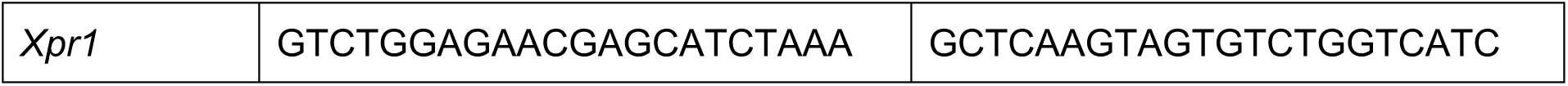

### Animals

All animal experiments adhered to the ARRIVE guidelines. The study involved male and female Balb/c mice or C57BL/6J mice (obtained from Jackson Laboratories). Mouse care followed institutional animal care and use guidelines, with approved experimental protocols from the Institutional Animal Care and Use Committee of the University of Michigan. Animals were housed at a density of 4 or 5 mice per cage under specific pathogen-free conditions. Daily monitoring was conducted throughout the experiments. All dissections were conducted aseptically using sterile equipment.

### Ex-Vivo Calvarial Cultures

Mouse calvarial explants were extracted from 24-day-old male Balb/c mice (Jackson Laboratories) and treated with increasing Pi and TGF-β2 protein concentrations. The culture procedure was adapted from^60^. Calvarial explants were grown in 96 well tissue culture-treated plates and maintained in an incubator at 37°C with 5% CO2, cultured in Minimum Essential Media α (MEM α) 1X with ascorbic acid (Gibco, Thermo-Fisher Scientific), supplemented with 10% Fetal Clone III (FCIII, HyClone), and 1X Antibiotic-Antimycotic (Invitrogen, Thermo-Fisher Scientific). Calvariae were pre-treated for one hour with 0.4ng/ml TGF-β2 or a vehicle control (4mM HCl + 0.1% BSA). Media was collected before being re-supplemented with respective concentrations of culture medium and treatment at experimental time points and calvaria punches were collected upon conclusion of the experiment; all samples collected and immediately frozen at -80°C for storage.

### α-TGFβ In Vivo Animal Study

To evaluate the endocrine response to high phosphate diets and anti-TGFβ antibody treatment, 11 week old C57BL/6J mice were placed on a high phosphate diet (1.8% Pi w/w, Teklad-Inotiv, TD.140507) or normal phosphate (0.6% Pi w/w, Teklad-Inotiv, TD.110362) then injected with an anti-TGFΒ antibody 1D11 or IgG control antibody at 5mg/kg three times over one week and then sacrificed. Serum, urine, and tissues were collected then flash frozen in liquid nitrogen and stored at -80°C. Bone marrow was flushed from long bones before being flash frozen and stored at -80°C.

### Tissue Homogenization

To protect against RNAse activity, we conducted RNA extraction from the mouse tissue at liquid nitrogen temperature. Tissue homogenization was conducted using a Bessman tissue pulverizer. Homogenized tissue was prepared for purification using TRIzol RNA Isolation Reagents (Invitrogen, ThermoFisher Scientific) as detailed previously.

### Free Mineral Concentration Quantification: Phosphate

Extracellular free phosphate concentrations were carried out with either the Phosphate Assay Kit (Colorimetric) (ab65622, Abcam) or the Malachite Green Phosphate Assay Kit (MAK307, Sigma-Aldrich). Absorbances were recorded and quantified using the Tecan Spark multimode microplate reader (Tecan Trading AG). All methods followed manufacturer protocol.

### Serum and Urine Chemistry

Serum was collected from mice and analyzed for multiple circulating factors. Intact bioactive FGF23 (iFGF23) was measured by ELISA (60-6800, QuidelOrtho). Additional ELISAs were used to quantify serum TGFβ2 (DY7346-05, R&D Systems), serum intact parathyroid hormone (iPTH) (60-2305, QuidelOrtho), and urine creatinine concentrations (0430, Stanbio Laboratory). Serum and urine phosphate levels were measured by colorimetric quantification (MAK307, Sigma-Aldrich). Absorbances were recorded and quantified using the Tecan Spark multimode microplate reader (Tecan Trading AG).

### Lead contact

Further information and requests for resources and reagents should be directed to and will be fulfilled by the lead contact, Lauren Surface.

## Materials, data and code availability

Material generated in this study, including MC3T3-E1 *Dmp1-2A-mCherry* cells will be made available to the field upon request. RNASeq and CRISPRi full sequencing data sets will be made available through GEO. ^1^

**References**

## Acknowledgements

We thank all members of the Surface and Mannstadt labs for helpful discussions, as well as the MGH Endocrine Unit, including Marc Wein, Petra Simic, Murat Bastepe, and Harald Jueppner. This study was supported by the University of Michigan Biological Sciences Scholar Program. L.E.S. is supported by NIH R00AR073903 (NIH/NIAMS), R56DE033668 (NIH/NIDCR), and a P30 core P&F award (AWD018648) from the Michigan Integrative Musculoskeletal Health Center (MIMHC) (supported by NIH/NCRR S10RR026475-01). We thank Paola Divieti Pajevic for generously providing the OCY454 cells used in this study. Fluorescence-activated cell sorting was performed at the Wellman Center for Photomedicine Core (MGH), Center for Regenerative Medicine Core (MGH), and the University of Michigan Medical School Flow Cytometry Core. Sequencing was performed by BGI Genomics (RNA-seq), MGH Next-Gen Sequencing Core (CRISPRi screen sequencing), and the University of Michigan Medical School Advanced Genomics Core (CRISPRi screen sequencing). Illustrations in Figure 4 were created in BioRender. Surface, L. (2025) https://BioRender.com/qof31y1

## Author Contributions

Conceptualization: E.K.Z, M.M., L.E.S

Investigation: E.K.Z., D.P.K, L.T. P.E.K., L.E.S

Writing: E.K.Z, L.E.S.

Funding Acquisition: M.M., L.E.S.

