## Supplemental Figures, Tables, and Figure Legends for "TGFβ signaling regulates the response of the skeleton to phosphate": Zhu_SuppFigs.pdf

**Supplementary Figure 1. Inorganic phosphate affects gene expression in osteogenic-lineage cell lines. a.** Gene expression as measured by RT-qPCR following exposure of undifferentiated MC3T3-1 and OCY454 cells to 4 or 10mM NaPi or 4 or 10mM $NaSO_4$ for the indicated lengths of time, normalized using *Rp*. Error bars represent standard deviation of technical triplicates, and indicative of at least n=3 separate experiments. **b.** Experimental design of NaPi/$NaSO_4$-treated cells for RNAseq. **c.** Heatmap of all differentially expressed genes (DEGs) from all the time points (30 minutes, 6 hours, 48 hours) in MC3T3-E1 cells exposed to 10mM NaPi vs. 10mM $NaSO_4$ (from DEseq2 with parameters $\log_2$(fold change) > |1| & adjusted p-value < 0.05)). **d.** Overlap of differentially expressed genes from all three timepoint in MC3T3-E1 and OCY454 cells reveals an overlap of 103 DEGs. **e.** Top gene ontology categories from Metascape analysis[1] of the 103 common DEGs from (D). **f.** Gene expression as measured by RNAseq of select DEGs after treatment with 10mM NaPi vs. 10mM $NaSO_4$, *: adjusted p-value < 0.05 (from DEseq2).

**Supplementary Figure 2. Phosphate-mediated induction of Dmp1-2A-mCherry and SEMA7A.**
MC3T3-E1 *Dmp1-2A-mCherry* cells are used for all experiments in this figure. **a.** Flow cytometry analysis of percent of mCherry+ cells following exposure to increasing NaPi over 2-4 days. n=6 biological replicates. **b.** Flow cytometry analysis of percent of SEMA7A+ cells following exposure to increasing NaPi for 12 hours – 2 days. **c.** Percent of mCherry+ cells following a 4 day exposure to 1,4, or 10 mM NaPi + 30 minute pre-treatment with vehicle, 10μM U0126 (MEK1/2 inhibitor), or 10nM 1,25 vitamin D. n=2-3 biological replicates. **d.** Percent of mCherry+ cells following 1 day incubation with 7mM $NaSO_4$, 4mM NaPi, or 7mM NaPi + 30 minute pre-treatment with vehicle, FGFR inhibitor (FGFRi, 100nM PD173074), 2.5mM $CaCl_2$, or 2.5mM $CaCl_2$ + FGFRi. **e.** Percent of mCherry+ and **f.** SEMA7A+ cells following differentiation for the indicated number of days in ascorbic acid and either 2mM or 10mM beta-glycerophosphate (βGP). **g.** MC3T3-E1 *Dmp1-2A-mCherry* were differentiated for 21 days with ascorbic acid and 2mM or 10mM βGP. During the last 3 days of differentiation, cells were exposed to 1mM, 4mM, or 10mM NaPi in addition to the ascorbic acid and βGP. The % of mCherry expressing cells was quantified via flow cytometry. Significance was assessed by two-way ANOVA, and corrected for multiple comparisons using Bonferroni's method (*: adjusted p-value<0.05, ns: not significant).

**Supplementary Figure 3.**
MC3T3-E1 *Dmp1-2A-mCherry* dCas9-KRAB cells are used for all experiments in this figure. **a.** Flow cytometry analysis to detect expression of CD81 and mCherry following lentiviral infection of vectors expressing either non-targeting or *Cd81*-targeting sgRNAs. **b.** Flow cytometry analysis of percent of mCherry+ cells following exposure to increasing 1mM or 10mM NaPi for 3 or 4 days following lentiviral infection of vectors expressing either non-targeting or *Dmp1*-targeting sgRNAs.

**Supplementary Figure 4.**
MC3T3-E1 *Dmp1-2A-mCherry* cells are used for all experiments in this figure. Cells were treated for 24 hours with either 4mM NaPi, 10mM NaPi, or 10mM NaSO4 with a 1 hour pre-treatment of either vehicle, TGF-β1 , TGF-β2, or TGF-β3 (at 0.4ng/ml). **a.** *Xpr1* gene expression was assessed by RT-qPCR, with fold change relative to cells treated with vehicle and 10mM $NaSO_4$, normalized using expression of *Rplp0*. n=2-3 technical replicates. ***b.*** *Slc20a2* gene expression was assessed by RT-qPCR, with fold change relative to cells treated with vehicle and 4mM $NaSO_4$, normalized using expression of *Rplp0*. n=3 biological replicates. ***c.*** *Enpp1,* and **d.** *Dmp1* gene expression was assessed by RT-qPCR, with fold change relative to cells treated with vehicle and 10mM $NaSO_4$, normalized using expression of *Rplp0*. n=2-3 technical

replicates. **e.** *Ltbp3* gene expression was assessed by RT-qPCR, with fold change relative to cells treated with vehicle and 10mM $NaSO_4$, normalized using expression of *Rplp0*. n=2-3 technical replicates. Significance was assessed by two-way ANOVA, and corrected for multiple comparisons using Bonferroni's method (*: adjusted p-value<0.05, ns: not significant).

**Supplementary Data 1:** Tables of normalized gene counts (fragments per kilobase per million reads, FPKM) and differentially expressed genes in MC3T3-E1 cells treated with 10mM NaPi or 10mM $NaSO_4$ for 30 minutes, 6 hours, or 48 hours.

**Supplementary Data 2:** Tables of normalized gene counts (fragments per kilobase per million reads, FPKM) and differentially expressed genes in OCY454 cells treated with 10mM NaPi or 10mM $NaSO_4$ for 30 minutes, 6 hours, or 24 hours.

**Supplementary Data 3:** Tables of MAGeCK analyses of the genome-wide CRISPRi screens for SEMA7A and Dmp1-2A-mCherry reporters in MC3T3-E1 cells.

1. Zhou, Y. *et al.* Metascape provides a biologist-oriented resource for the analysis of systems-level datasets. *Nat. Commun.* **10**, 1523 (2019).

### Supplementary Figure 1

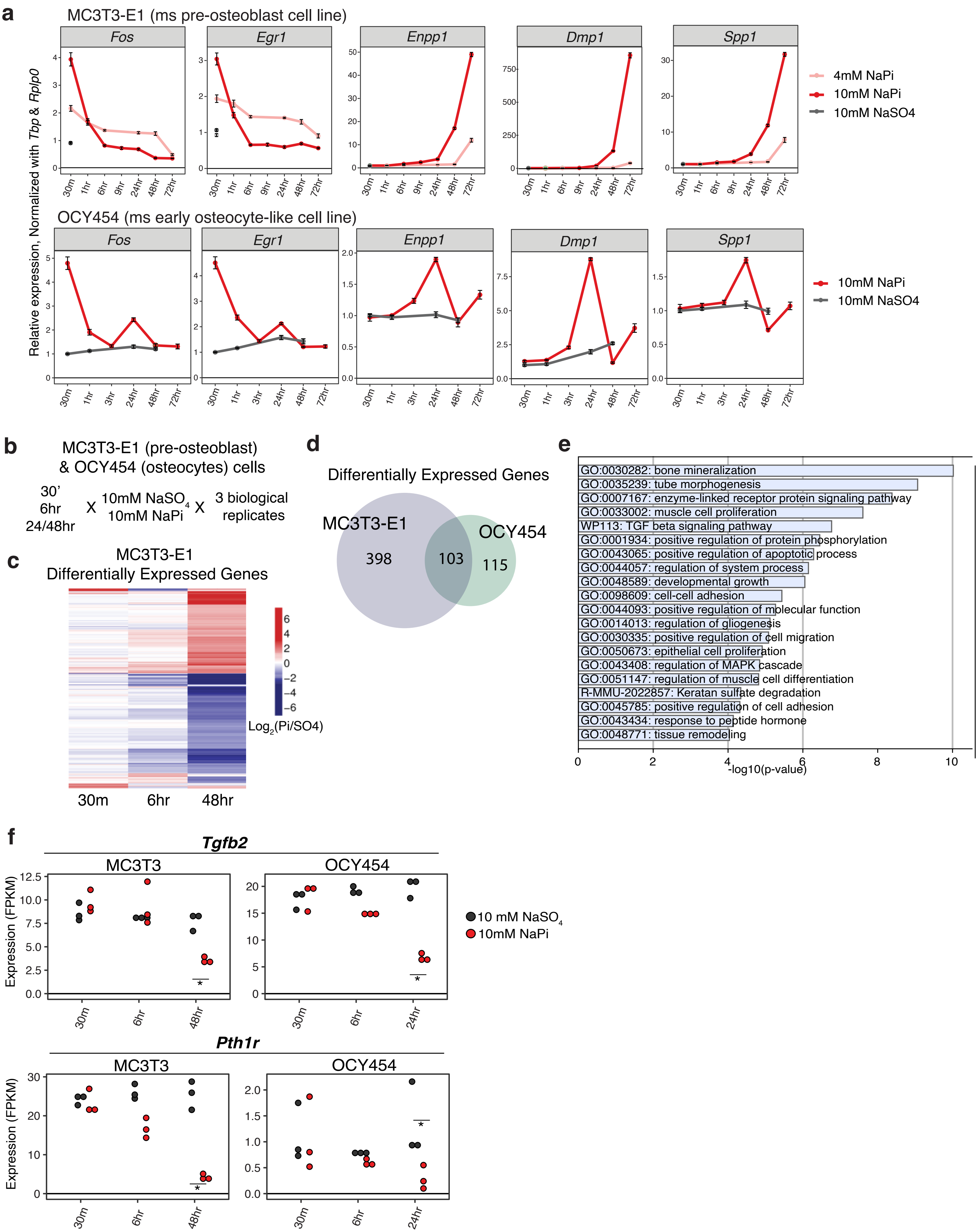

### Supplementary Figure 2

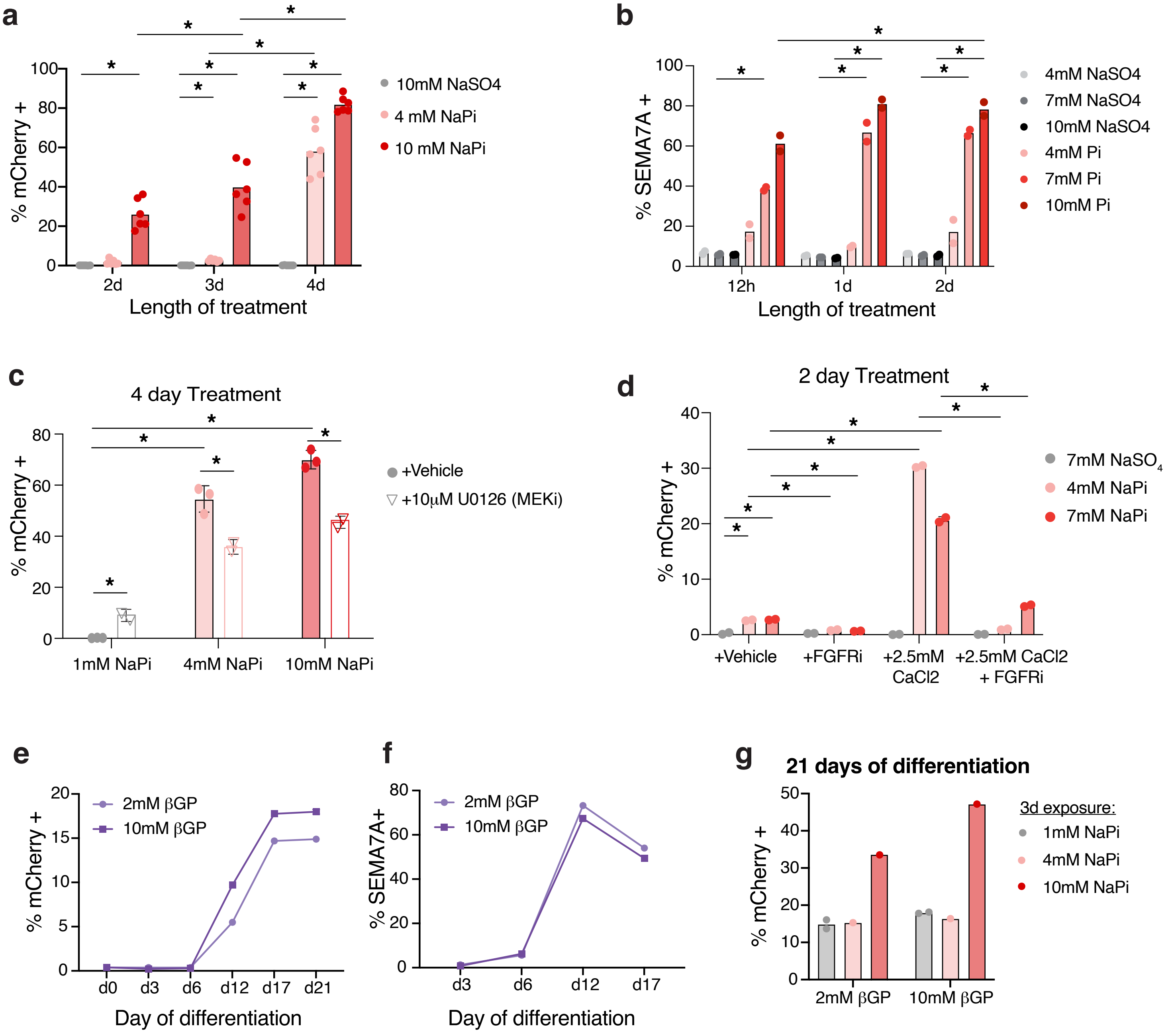

### Supplementary Figure 3

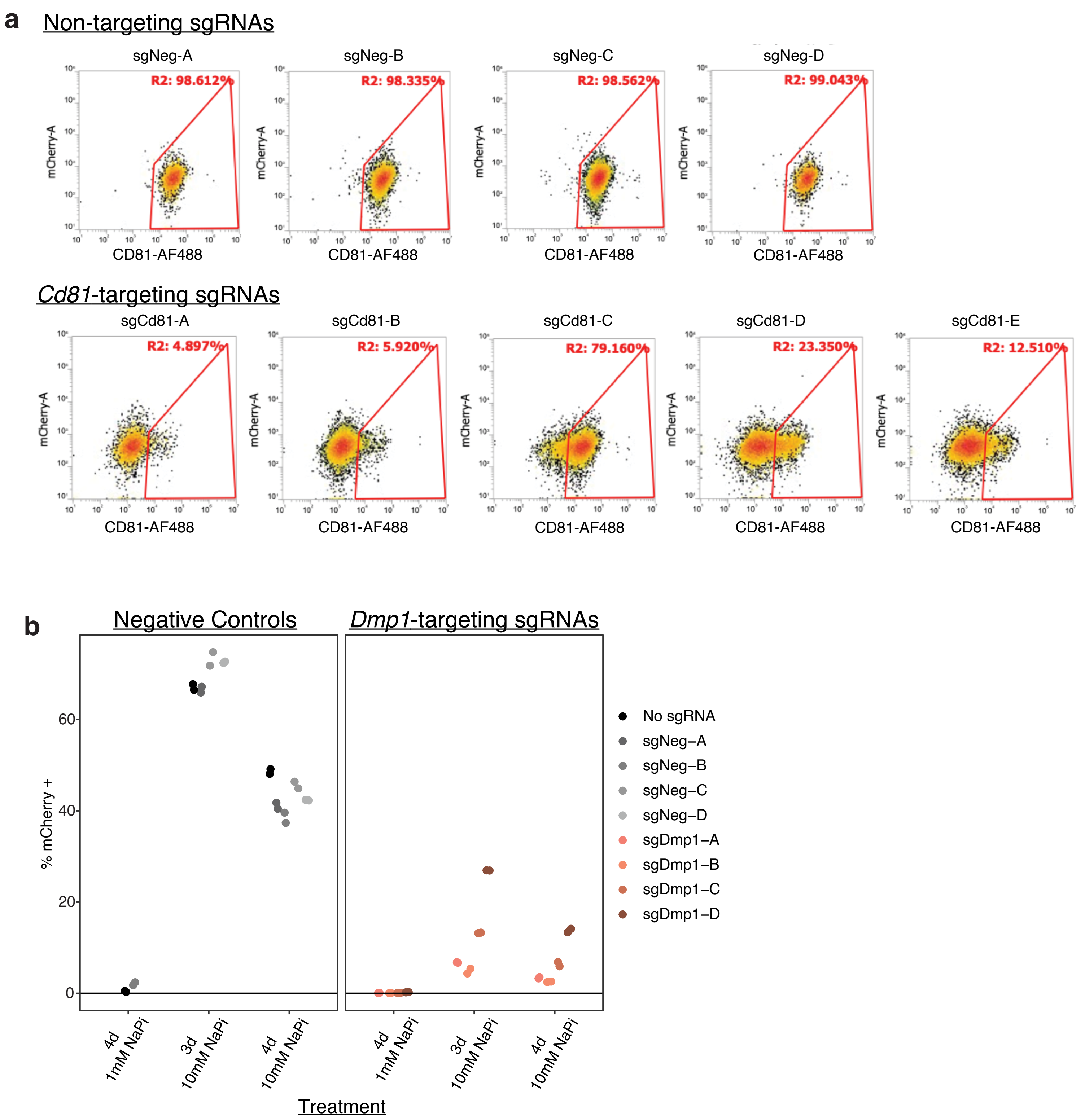

### Supplementary Figure 4

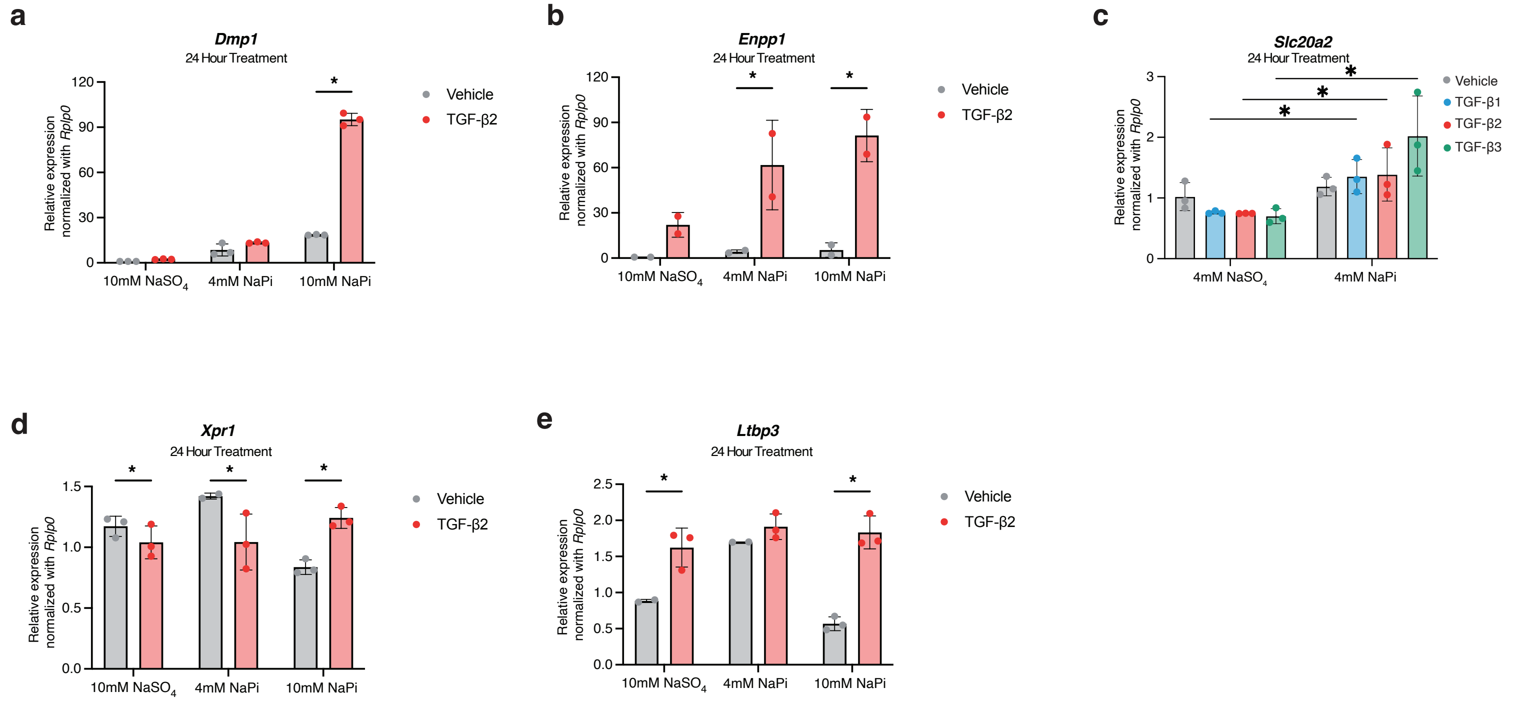
